# A transgenic mouse allows to capture the HLA-C*06:02 immunopeptidome in a model of psoriasis

**DOI:** 10.1101/2025.05.01.651638

**Authors:** Asolina Braun, Jesse I. Mobbs, Shanzou Chung, Johanna E. Tuomisto, Chen Li, Katherine E. Scull, Rochelle Ayala, Sushma Anand, Sri Ramarathinam, Johannes S. Kern, Anne Fourie, Murray McKinnon, Carl Manthey, Daniel G. Baker, Navin Rao, Nicole A. Mifsud, Patricia T. Illing, Julian P. Vivian, Jamie Rossjohn, Anthony W. Purcell

**Author notes:** Correspondence; (contributed equally).

## Abstract

Psoriasis vulgaris is a T cell-mediated autoimmune skin condition affecting around one in fifty people worldwide. Whilst advanced immunomodulatory therapeutic options have become available in recent years, ongoing disease suppression is still required with no curative treatment available to date. The human leukocyte antigen class I allele HLA-C*06:02 is the main genetic risk determinant of psoriasis. Its function is to present peptide antigens to CD8^+^ T cells and natural killer cells which in turn elicit and perpetuate the immune response, yet little is known about ligands presented by HLA-C*06:02. To gain an understanding which HLA-C*06:02-restricted peptides are presented by epidermal cell populations and might be initiators of the autoimmune response in psoriasis, we have conducted an in depth immunopeptidomics analysis of HLA-C*06:02^+^ keratinocyte and melanocyte cell lines. Furthermore, we introduce a HLA-C*06:02 transgenic mouse which, in conjunction with the imiquimod model of psoriasis, allowed us to assess the *ex vivo* immunopeptidome of HLA-C*06:02 in psoriasiform skin. Overall, we detected 20,812 high confidence HLA-C*06:02 bound peptide ligands. Thus, we present a comprehensive coverage HLA-C *in vitro* immunopeptidomics dataset and the first HLA-C*06:02 *ex vivo* dataset of psoriasis-relevant peptide antigens that may inform the development of antigen-specific, novel curative therapeutic approaches in psoriasis.

## Introduction

The carriage of the human leukocyte antigen (HLA)-C*06:02 allotype has consistently been recognised across different global populations as the major genetic risk for developing psoriasis ^1-3^. HLA molecules, encoded within the highly polymorphic major histocompatibility complex (MHC), function to present peptide antigens to immune cells. HLA-C*06:02 belongs to the MHC class I type (HLA I) and displays peptides derived from intracellular sources on the cell surface. HLA-C*06:02 is a common and well-documented HLA allele with a global frequency of around 6% (pypop.org). However, in psoriasis patients it is far more frequent, with a prevalence of 50% overall and >80% in early onset psoriasis cohorts^4^.

HLA-C is recognised not only by CD8^+^ T cells via their T cell receptor (TCR), but can also be recognised by the killer cell Ig-like receptor (KIR) expressed on natural killer (NK) cells, effector T cells and NKT cells. This dual role of HLA-C as an activating and inhibitory molecule is the driver of the tightly regulated expression of HLA-C on the cell surface^5-7^. The low constitutive levels of HLA-C expression also likely explains the current lack of a dataset of HLA-C*06:02-presented epidermal ligands. Notably, in healthy and allergic eczema skin, HLA-C expression is more diffuse and higher in the epidermis than dermis, however histological staining of psoriatic lesions indicates HLA-C*06:02 is more highly expressed in the basal epidermis which harbours highly proliferative basal keratinocytes, Langerhans cells and melanocytes^8^.

The study of MHC ligands, also known as immunopeptidomics, involves the isolation of peptide-MHC complexes via immunoaffinity chromatography, the dissociation of the MHC-bound peptides which are subsequently separated via reversed-phase high-performance liquid chromatography (HPLC) or filtration and analysed using liquid chromatography-tandem mass spectrometry (LC-MS/MS)^9^. By using this approach in a B lymphoblastoid cell line, we have previously determined the HLA-C*06:02 peptide binding motif which features arginine residues at anchor positions P2 and P7 and hydrophobic amino acids at P9^10^. Now we apply immunopeptidomics to interrogate psoriasis-relevant HLA-C*06:02-restricted peptide ligands in more physiologically relevant settings, including epidermal cell lines and a HLA-C*06:02 transgenic mouse to study antigen presentation in psoriasiform skin directly *ex vivo*. This approach has generated a comprehensive database of 20,812 HLA-C*06:02 ligands that are of high relevance for antigen discovery of psoriasis molecular triggers in genetically susceptible patients.

## Results

### HLA-peptide identification using the immunopeptidomics workflow on epidermal cells

To determine the peptide repertoire of epidermal cell populations, we used the 1106 keratinocyte and VMM1 melanocyte cell lines that naturally express the HLA-C*06:02 allotype. The cells were either used directly or pre-treated with IFN-γ to investigate peptide antigens presented in steady state or under inflammatory conditions. IFN-γ is one of the cytokines upregulated in psoriatic plaques, it is known to upregulate HLA expression and to induce the formation of the immunoproteasome thereby remoulding the immunopeptidome^11-13^. Flow cytometric analysis of HLA-C or pan-HLA class I expression on the cells at the collection timepoint confirms that IFN-γ upregulated HLA I in general as well as HLA-C specifically (**Fig. 1A**). Furthermore, we have transformed the keratinocyte cell line A431 to express soluble HLA-C*06:02 (A431_sC6) to overcome physiologically low HLA-C*06:02 expression levels observed in some cell lines. Expression of soluble HLA has the advantages of combining relatively high yields of HLA-C with the ease of processing soluble HLA-peptide complexes from the cell culture supernatant that recapitulate the physiological array of bound peptide ligands. This has been previously validated by the unaltered consensus binding motif of peptides isolated from soluble HLA-secreting cell lines, including 721.221 cells expressing soluble HLA-C*06:02^10, 14^. To generate the HLA-C*06:02-restricted, epidermal cell-derived immunopeptidome we followed protocols that we have established and described in detail previously^9^. In short, cell lines were expanded, in part treated with IFN-γ for 48h, cell pellets (for membrane HLA) or culture supernatants (for soluble HLA) were collected and peptide-HLA complexes captured by immunoaffinity purification. HLA-C was isolated using the antibody DT9^15^. Upon acid elution of peptide-HLA complexes, peptides were separated by reverse-phase HPLC, identified by LC-MS/MS and analysed using the PEAKS Online software suite with a false discovery rate cut-off of 5 percent used to qualify peptide identifications. Overall, we detected 33,525 non-redundant, DT9-eluted 8-12-mer peptides across all analysed epidermal cell lines, the individual detected numbers per cell line and condition are depicted in **Fig. 1B** and can be found in **Suppl. File 1**. In all cell lines, the length distribution featured predominantly 9-mers (**Fig. 1C**) as is expected for HLA class I presented peptides. To identify common and individually presented peptides, we have subjected the immunopeptidome dataset to Venn and Upset plot analysis (**Fig. 1D**). The top seven sets of peptides depicted in the Upset plot are either condition-specific peptides or cell line-specific peptides followed by various degrees of overlapping datasets between cell lines and conditions.

**Figure 1.**
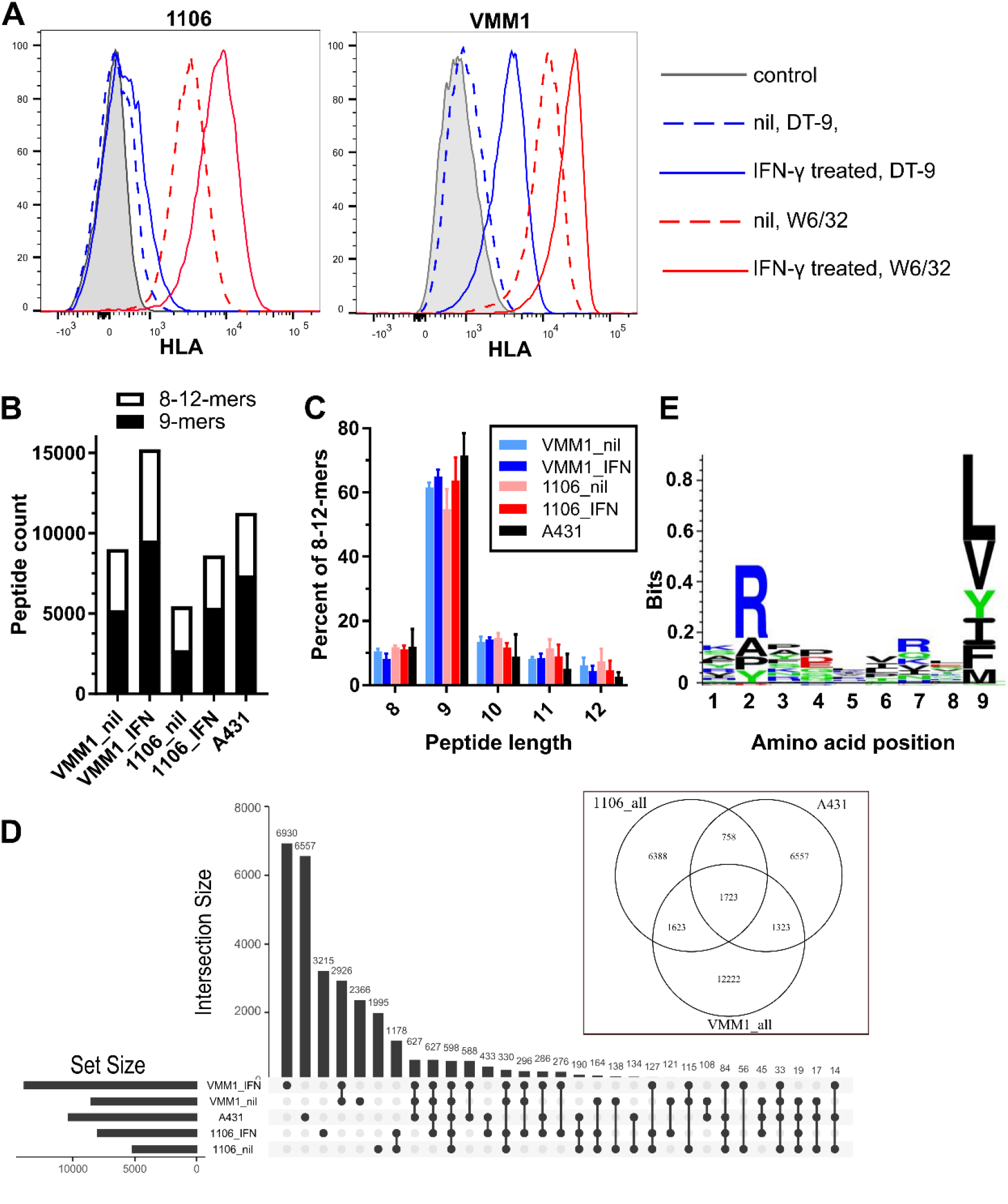
Characteristics of HLA-C ligands isolated from epidermal cell lines. (**A**) 1106 keratinocytes and VMM1 melanocytes were treated with IFN-γ (2 ng/ml, 48h) or left untreated before flow cytometric staining with the HLA-C specific DT9 antibody or W6/32 which stains HLA-A/B/C. Secondary detection antibody staining alone is included as a negative control; n= >3 independent experiments. (**B**) Cell pellets (VMM1, 1106) or supernatant of unstimulated A431 expressing soluble HLA-C*06:02 were subjected to immunoaffinity purification of HLA-C bound peptides and analysed via mass spectrometry. Detected numbers of 8-12-mer and 9-mer peptides are shown; n=3 independent experiments. (**C**) Length distribution of HLA-C bound peptides (8-12-mers shown). (**D**) Upset plot depicting common and unique peptide groups across all conditions. The Venn diagram shows overlaps between cell lines (peptides from untreated and IFN-γ-treated conditions per cell line are combined). (**E**) Sequence logo of DT9-isolated 9-mers from all three cell lines combined, generated after Gibbs clustering and filtering as per Suppl. Fig. 1. Heights of amino acids are proportional to their frequency of occurrence.

After removal of post-translational modifications (PTMs) and resulting duplicates, sequence logo analysis of 16,867 9-mers revealed a HLA-C*06:02 motif which was consistent with our and other previously published data^10, 16^ (**Suppl. Fig. 1A**). Thus, in line with previous reports of the HLA-C*06:02 immunopeptidome derived from B lymphoblastoid cell lines 721.221 and C1R, this immunopeptidome features a motif preference for arginine anchor residues at P2 and/or P7 as well as predominantly hydrophobic amino acids at the C-terminal (PΩ) position (**Figure 1E**)^10, 16^. A closer examination of the peptide motif in **Suppl. Fig. 1A** reveals some unexpected amino acids such as valine, proline and glutamic acid (V, P and E) at the key anchor position P2 and lysine (K) at the PΩ position. We hypothesised that this likely stems from peptides bound to co-isolated HLA-A and -B alleles, which are present at much higher levels on the cell surface than the targeted HLA-C molecules. To further assess this possibility, we performed Gibbs cluster analysis to extract sub-groups of peptides with similar characteristics. The HLA-C motif was always dominant, but in addition we found some smaller peptide clusters in each cell line (**Suppl. Fig. 1B, top row**). Subdominant clusters detected by Gibbs clustering displayed motifs that could be readily allocated to co-expressed HLA-A or -B allotypes (**Table 1 and Suppl. Fig. 1B, top row**). We next proceeded with the filtering of peptides resulting from the Gibbs cluster analysis. All peptides belonging to Gibbs clusters in line with the known HLA-C*06:02 motif and peptides not allocated to any cluster were retained. For any peptides allocated to a HLA-A/B cluster, we extracted peptides that were predicted to be HLA-C*06:02 binders (NetMHCpan4 rank of <2) as well as any 9-mers that contained an arginine at position 2 and/or 7 (HLA-C*06:02 anchors) under exclusion of peptides with a C-terminal lysine or arginine (likely derived from HLA-A*03:01/*11:01). This process resulted in a refined list of 13,289 9-mers and overall 19,779 8-12-mer HLA-C*06:02 ligands (marked as “high confidence” in **Suppl. File 1**). The peptide motif of these refined 9-mer peptides had a markedly lower representation of valine, proline and glutamic acid on P2 and lacked an overrepresentation of lysine on PΩ (**Fig. 1E**; also in **Suppl. Fig. 1** compare **B bottom left** to **A**). It needs to be noted that all three cell lines also expressed HLA-C*07:02. Whilst HLA-C*06:02 features two arginine anchors at P2 and P7 at similar frequencies, HLA-C*07:02 preferably presents peptides featuring an arginine at P2. Given that HLA-C*06:02 and HLA-C*07:02 present very similar peptides, it is not possible to distinguish whether peptides with a P2 arginine anchor are derived from either of these HLA-C alleles. In all likelihood, peptides bound to HLA-C*07:02 can also be presented by HLA-C*06:02 (as is assessable by affinity binding prediction) and thus were retained in a final list of a range of diverse and unique peptide groups across three epidermal cell lines under steady state and pro-inflammatory conditions.

**Suppl. Fig. 1.**
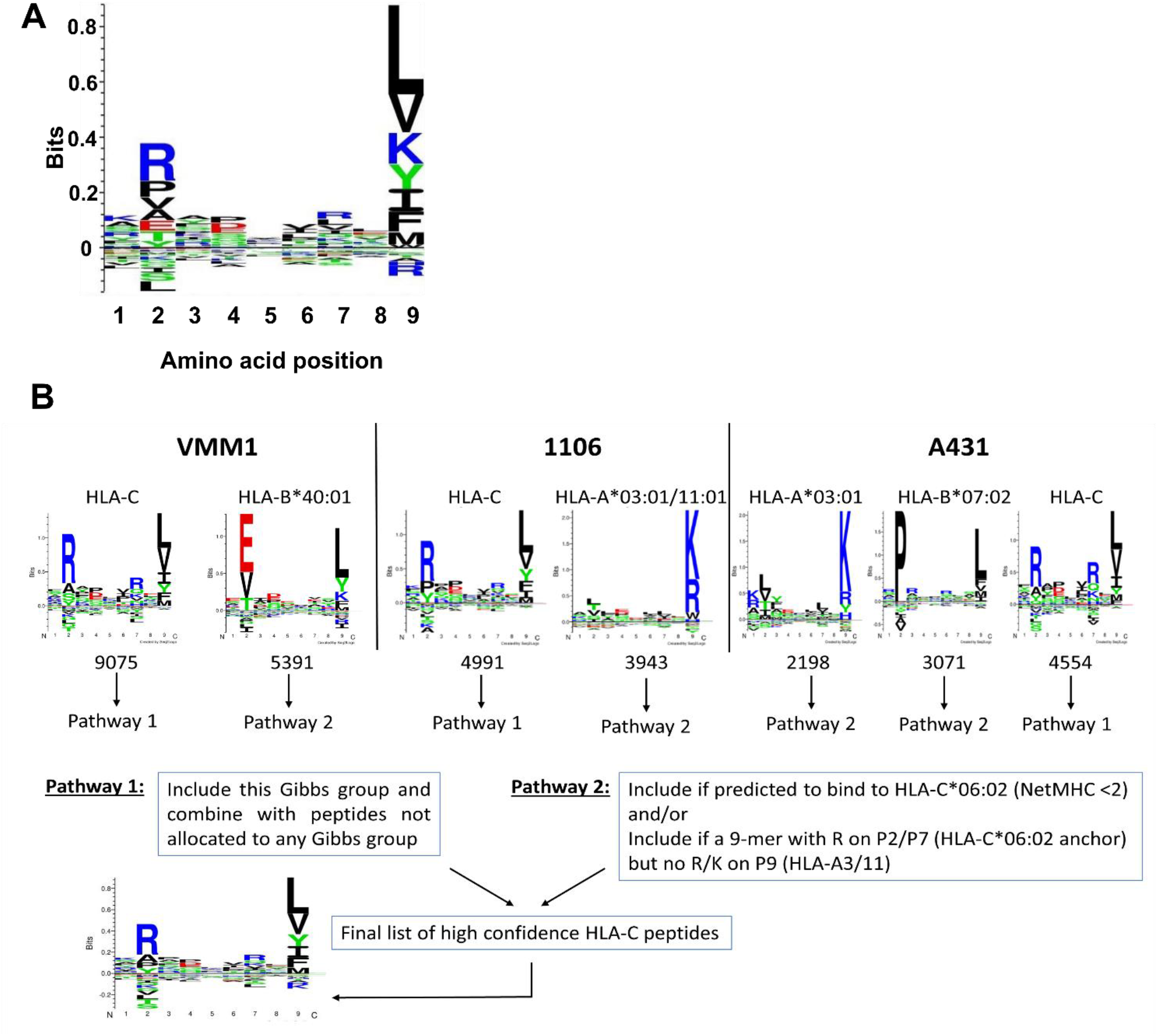
Filtering of HLA-C ligands isolated from epidermal cell lines. The Gibbs clustering approach was used to detect distinct peptide groups. Peptides allocated to a HLA-C motif cluster were included as well as the remaining peptides that were not specifically allocated to any Gibbs cluster (Pathway 1). Peptides from HLA-A/B cluster groups were further filtered to align with HLA-C*06:02 predicted binding and its known motif (Pathway 2). A total of 19,779 high confidence HLA-C*06:02 peptides was retained after filtering.

**Table 1.** Haplotypes of cell lines used in this study.

| Cell type | Cell line | HLA-A | HLA-B | HLA-C | HLA II |
| --- | --- | --- | --- | --- | --- |
| Keratinocyte | 1106 | 03:01<br>11:01 | 07:02/22<br>57:01 | 06:02<br>07:02/234 | DRB1*01:01, DRB1*07:01<br>DPB1*04:01<br>DQB1*03:03, DQB1*05:01<br>DQA1*01:01/4/5, DQA1*02:01 |
| Keratinocyte | A431 | 03:01 | 07:02/22 | 07:02/27/234 | DRB1*11:04<br>DPB1*15:01<br>DQB1*03:01<br>DQA1*05:05/19 |
| Melanocyte | VMM1 | 03:01:01G<br>26:01:01G | 40:01:01G<br>51:01:01G | 06:02<br>07:01 | DRB1*13:02, DRB1*15:01<br>DPB1*02:01, DPB1*04:01<br>DQB1*06:02, DQB1*06:04<br>DQA1*01:02 |

Next, we performed ontogeny/pathway analysis on the source antigens. When compared to the previously published HLA-C*06:02 ligands from B cell lines^10, 16^, this dataset was enriched for developmental processes, anatomical structures and cell junction organisation (**Suppl. File 2**, sheet GOrilla_1). To further understand which skin-related biological processes were represented in our data, we compared our immunopeptidome against the 602 skin-elevated genes in the Human Protein Atlas and found that we could detect 118 of them in our dataset^17^. When comparing these 118 skin-elevated genes to the entire immunopeptidome dataset, we found that among others, the highly enriched processes were cornification, epidermis development, keratinisation, cell-cell adhesion, hemidesmosome assembly, hair cycle and skin barrier establishment (**Suppl. File 2**, sheet GOrilla_2). The broad representation of dermatological processes in our dataset is of high relevance. The identification of differentially expressed genes and proteins in numerous transcriptomic and proteomic studies of psoriasis lesional biopsies to date is primarily a reflection of the ongoing structural changes, inflammation and the influx of immune cells. Here however, the immunopeptidome dataset now allows us to explore which tissue-associated peptide antigens might be available for recognition to immune cells during the initiation, development and perpetuation of psoriatic lesions in high-risk HLA-C*06:02^+^ patients.

### Generation of transgenic mice to study the *ex vivo* skin-derived HLA-C*06:02 immunopeptidome

After capturing HLA-C*06:02-bound ligands from cell lines, we wanted to investigate the presentation of HLA-C*06:02-derived peptides in skin directly *ex vivo*. We have developed a HLA-C*06:02 transgenic mouse (HLA-Cw6^Tg^), expressing the tandem HLA-C*06:02 transgene and human beta-2-microglobulin (β2M) as a single chain construct under the ubiquitous and physiological mouse β2m promoter on a C57/BL6 mouse background. Phenotyping of various human (hu) and murine (mu) components of the antigen presentation pathway (hu-β2M, mu-β2m, mu-MHCI) by flow cytometry revealed distinct expression levels commensurate with genotype (**Fig 2A**). HLA-Cw6^homo^ mice lacked mu-β2m expression as expected, while HLA-Cw6^het^ mice displayed very low expression levels of mu-β2m. The opposite was true for hu-β2M expression levels, HLA-Cw6^homo^ showed the highest level of expression, expression in HLA-Cw6^het^ was slightly lower, yet markedly higher than wildtype mice. Notably, some cross-reactivity of the anti-mu-MHCI antibody to HLA-C*06:02 could be observed. We next proceeded to evaluate the expression levels of HLA-C*06:02 in transgenic mice. Due to the intrinsically low expression of HLA-C*06:02 under the physiological β2m promoter, we looked at the expression under inflammatory conditions.

**Figure 2.**
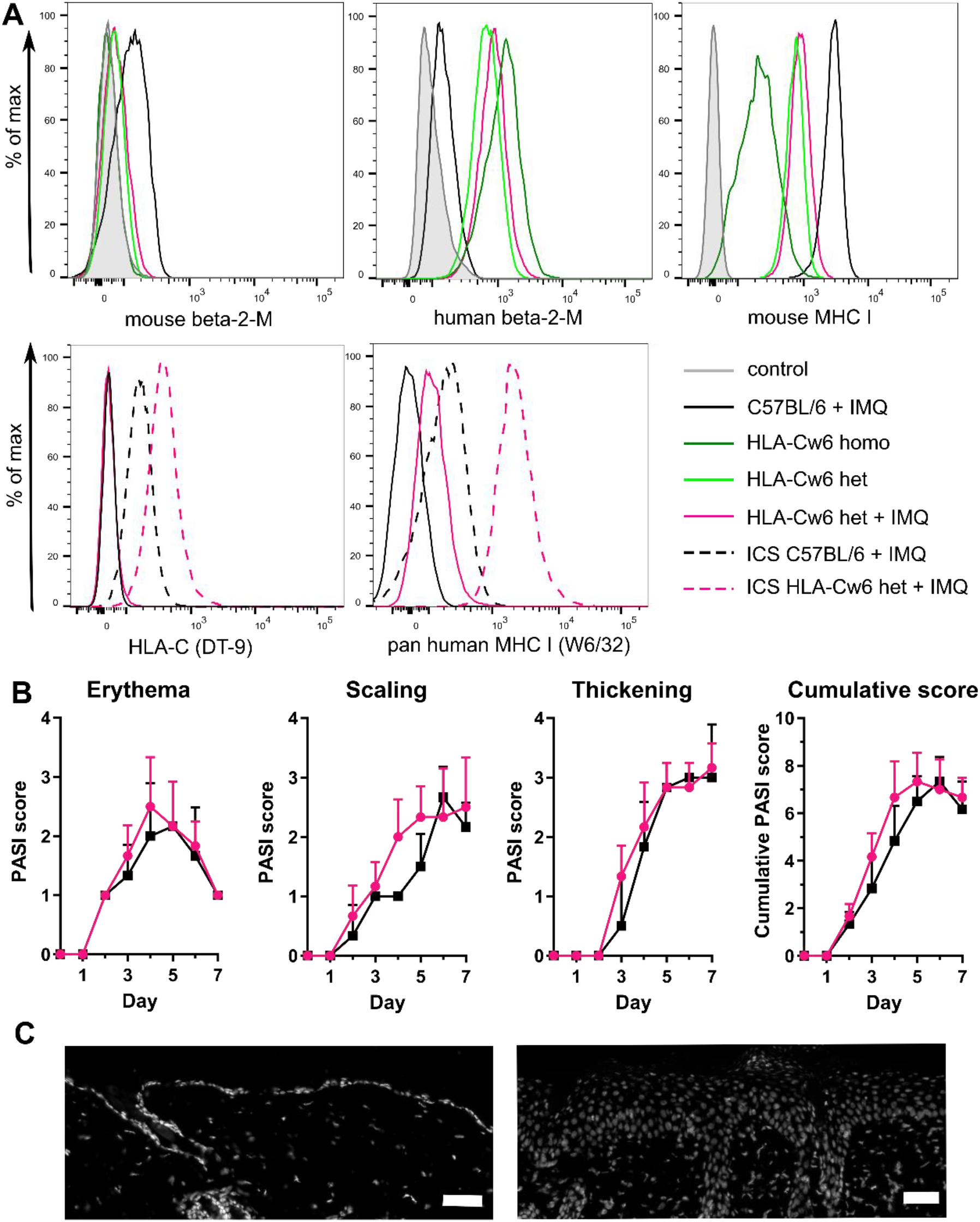
Phenotyping of the new HLA-C*06:02Tg mouse strain. (A) Cell surface expression or intracellular staining (ICS, dashed lines) of murine and human beta-2-microglobulin, mouse MHC class I and human HLA-C on splenic T cells of untreated mice or mice after eight consecutive topical applications of imiquimod (IMQ). Minimum three independent experiments. (B) Representative skin PASI scoring of wildtype (black) and HLA-C*06:02het (pink) IMQ-treated mice, SD is shown. (C) Representative DAPI staining of skin from untreated mice (left) or skin of mice treated with IMQ (right), bar=50 µm. Data collectively from one of three independent experiments with 5 mice per IMQ-treatment group.

Topical application of imiquimod (IMQ, Aldara™), a TLR7/8 agonist, is a well-established mouse model of psoriasis and is known to elicit not only a local, but also a systemic inflammatory response^18^. To investigate HLA-C*06:02 expression levels in this inflammatory condition, we analysed splenocytes from wildtype and HLA-Cw6^het^ mice upon 8 consecutive daily topical applications of IMQ. Heterozygous mice were chosen to overcome any potential limitations connected to the requirement of β2m in the formation of the neonatal Fc receptor which performs important functions in immunoglobulin and albumin binding as well as antigen presentation^19^. We included the HLA-C specific DT9 and the high affinity anti-pan human MHC class I antibody W6/32 in the analysis. Surface staining with DT9 was negligible while W6/32 was able to detect a low-level expression of HLA-C*06:02 on transgenic splenocytes (**Fig. 2A**). Given that both antibodies could readily detect HLA-C*06:02 intracellularly (dashed lines), the low expression of HLA-C*06:02 on the cell surface was likely due to physiological regulation which is known to tightly control HLA-C membrane transport^5, 6^. Furthermore, we assessed the skin inflammation induced by IMQ and found that HLA-Cw6^het^ and wildtype mice exhibited similar erythema, scaling, skin thickening and overall cumulative psoriasis area and severity index (PASI) scores (**Fig. 2B**). Histology further confirmed acanthosis, a hallmark of psoriasis, in IMQ-treated skin (**Fig. 2C**).

### *Ex vivo* immunopeptidomics of psoriasiform skin

After having phenotyped and confirmed that we can detect membrane HLA-C*06:02 expression in transgenic mice via flow cytometry, we went on and performed immunopeptidomic analysis of psoriasiform skin, contralateral non-lesional skin and spleens of IMQ-treated HLA-Cw6^het^ and wildtype C57BL/6 mice. Based on flow cytometric analysis, we used the higher affinity pan-HLA class I W6/32 antibody rather than DT9 to immunoprecipitated HLA-C complexes, followed by the Y-3 antibody (anti-H-2K^b^). The bound peptide ligands were extracted and analysed by LC-MS/MS. After removing any peptides detected in wildtype mice, we retained 4004 non-redundant 8-12-mers with the vast majority of them (99.1%) exclusively detected in the W6/32, but not in the Y-3 elution from IMQ-treated HLA-Cw6^het^ mice, indicating a near-complete capture of HLA-C by W6/32 in the first step and minimal cross-reactivity of Y-3 with HLA-C. Overall, we detected 2570 8-12-mers from spleen, 1790 in psoriasiform skin and 1201 in non-lesional skin (**Fig. 3A**). The extracted peptides were mostly 9-mers and displayed a peptide motif that conformed with HLA-C*06:02-restricted human peptides, thus confirming the suitability of the new mouse model to study *in vivo* antigen presentation by HLA-C*06:02 (**Fig. 3A-C**). Next, we compared eluted 9-mer peptides from HLA-Cw6^het^ mice to the database of 19,779 PTM-stripped peptides previously isolated from human HLA-C*06:02^+^ cell lines. We found a high overlap: 45% of the murine 9-mers were either a perfect match (36%) or matched with only one amino acid difference (9%) to human-derived HLA-C*06:02-restricted ligands. Further analysis showed that 1,384 of 4,004 peptides were detected exclusively in murine HLA-C*06:02^Tg^ skin, not in their spleen, and 929 of these peptides were exclusive to (761/54%) or more abundant (168/12%) in psoriasiform skin. A closer examination of the 761 peptides uniquely detected in psoriasiform skin (**Fig. 3D** and **Suppl. File 3**, PsO unique sheet) revealed that 334 peptides (45%) were 9-mers and 256 (34%) were predicted to be HLA-C*06:02 binders. GOrilla pathway analysis of their source proteins showed an enrichment for intracellular components (**Suppl. File 3**, GOrilla sheet). Among the 14 enriched intracellular genes was Ezrin, a previously proposed potential target of streptococcal-induced autoimmune responses in psoriasis^20^. Indeed, in a recent systematic review of proteomic studies, Ezrin has been consistently found to be upregulated in patient lesional skin^21^. In order to further investigate the hypothesis of molecular mimicry in psoriasis, we compared the peptides uniquely detected in psoriasiform skin to an in-silico dataset of HLA-C*06:02 predicted binders from *Streptococcus pyogenes* strains and their human molecular mimics. We could find 14 mimics between streptococcal and human peptides where a 100% match to the human peptide was found in our murine dataset (**Suppl. File 3**, mimicry sheet). Among the streptococcal mimic peptides were five enzymes, transporters, repair proteins, surface proteins and the C5a peptidase virulence factor. Among the human mimic peptides, which were all found to be derived from proteins with expression in skin, were proteins involved in transcriptional regulation, chaperones as well as the known, skin-expressed structural proteins vimentin and the collagen type IV alpha 2 chain. Overall, we conclude that HLA-Cw6^Tg^ mice, in conjunction with the IMQ model of skin disease provide a new and highly useful *in vivo* model system to study antigen presentation in psoriasis.

**Figure 3.**
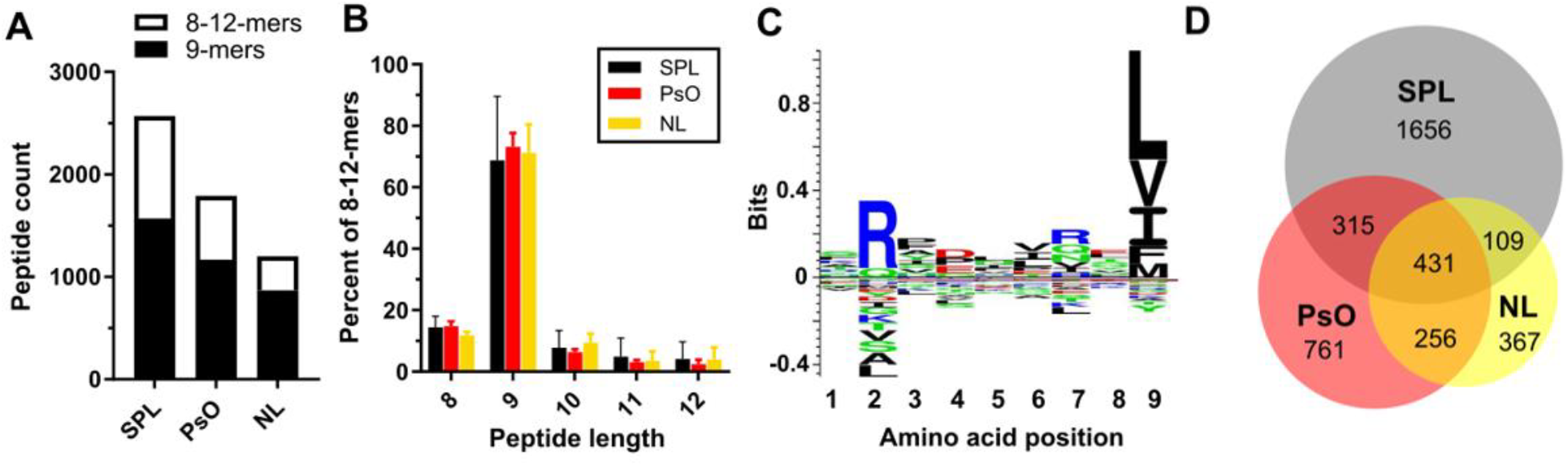
Immunopeptidome of HLA-C*06:02^Tg^ mice subjected to the mouse model of psoriasis. Spleens (SPL), psoriasiform skin (PsO) and non-lesional contralateral skin (NL) from 5 mice/group were pooled and HLA-C*06:02 ligands isolated using W6/32 antibody and analysed via LC-MS/MS. Any peptides also detected in C57BL/6 control mice were excluded from analysis. (A) Numbers of 9-mers and total 8-12-mers isolated per organ are shown. (B) Similar length distribution is observed across different organs. (C) Motif of 2340 9-mer peptides from the pool of the total 4004 8-12-mers detected. (D) Unique and redundant peptides identified across tissues. Pooled (A, C, D) or individual (B) data from 3 independent experiments is shown.

### Post-translational modification of peptide ligands

Specific PTMs have been previously associated with other autoimmune conditions and the breakdown of tolerance. For example, some insulin epitopes are posttranslationally modified in Type 1 Diabetes ^22, 23^. Similarly, a number of citrullinated epitopes elicit T cell and B cell responses in rheumatoid arthritis and treatment with tolerogenic dendritic cells exposed to citrullinated epitopes has been shown to have beneficial effects for patients in early stage clinical trials^24, 25^. Therefore, we extracted the most prevalent PTMs found in the immunopeptidome of epidermal cell lines and the *ex vivo* immunopeptidome of HLA-C*06:02^Tg^ mice. While methionine oxidation dominates the PTM landscape in cell lines, the PTM profile of peptides isolated *ex vivo* has a higher diversity, additionally featuring relatively high proportions of acetylated, amidated and deamidated peptides (**Suppl. Fig. 2**). It therefore appears that the *ex vivo* isolated immunopeptidome extends upon the *in vitro* data with the ability to detect a wider range of posttranslationally modified peptide antigens. This may be important in psoriasis, where deamidation could influence the integrity of epithelial barriers. In celiac disease, deamidation of gliadin peptides by tissue transglutaminase enhances their immunogenicity. This process can disrupt the epithelial barrier, leading to inflammation and immune activation^26^.

**Suppl. Fig. 2.**
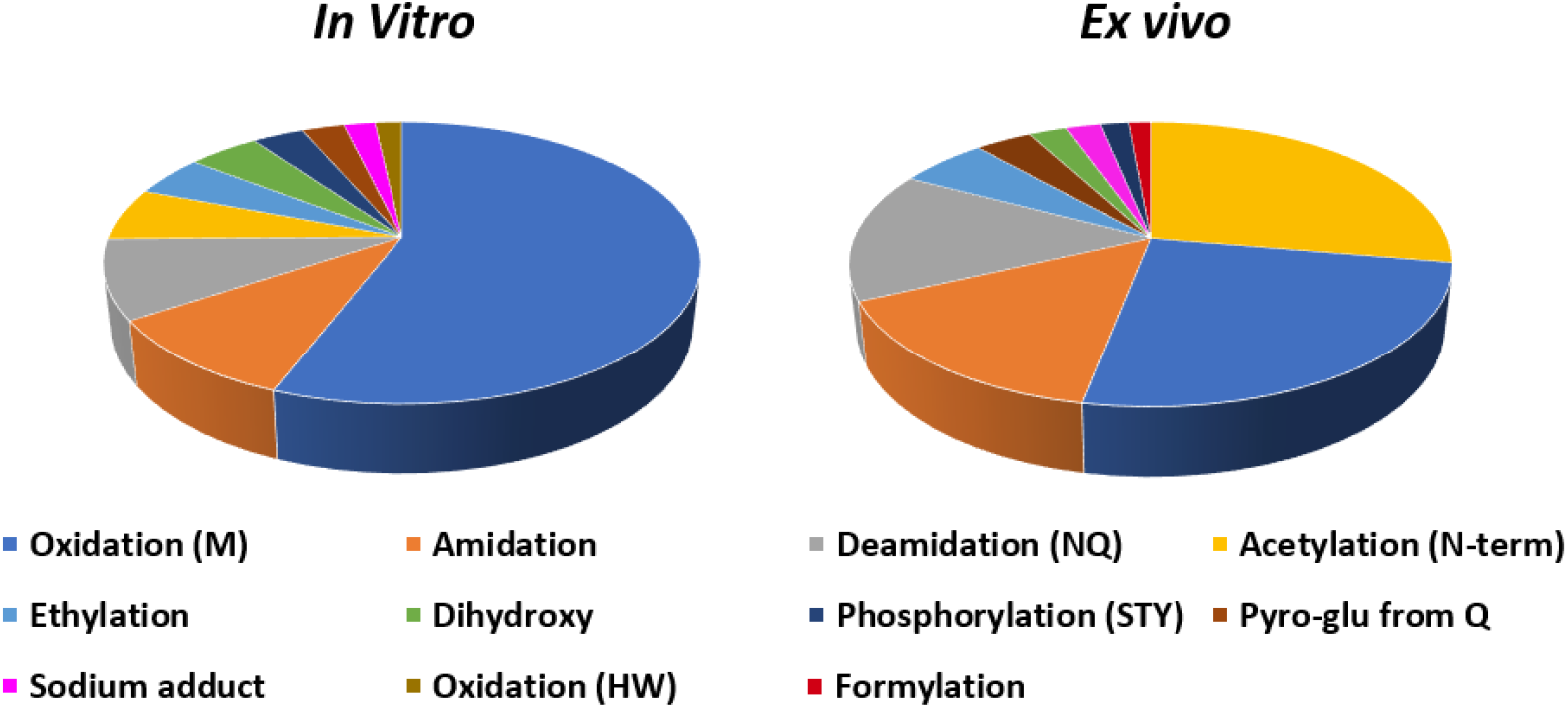
Analysis of PTMs in immunopeptidome datasets. PTM analysis was performed using the unrestricted PTM search in PEAKS. The top 10 PTMs from the 19,779 high confidence HLA-C6 derived peptides from *in vitro*-generated human cell lines (left) and the 4,004 peptides of the *ex vivo* murine immunopeptidome (right) were extracted (under the exclusion of carbamidomethylation which is not relevant to this project) and are shown in a pie chart with each PTM depicted as a portion of the top 10 PTMs.

## Discussion

Immunopeptidomics is a powerful approach enabling the investigation of antigen presentation by MHC molecules. Nevertheless, existing data on the immunopeptidome of HLA-C allotypes, especially in cells other than transgenic model B cell lines, is scarce. Here we report a comprehensive HLA-C*06:02 immunopeptidome derived from *in vitro* cultured epidermal cell lines. This newly generated dataset is enriched for peptides derived from skin-relevant proteins, allowing to explore new psoriasis trigger candidates and to interrogate skin-expressed peptides and proteins as triggers of psoriasis.

One suggested HLA-C*06:02-restricted trigger of psoriasis is the ADAMTLS5-derived peptide^27^. However, we could not detect the ADAMTSL5-derived proposed psoriasis trigger candidate peptide VRSRRCLRL in any of the present data, nor in HLA class I elutions of MEL33 and MEL45 melanocyte lines, both expressing HLA-C*06:02 (Aranha et al., manuscript in preparation). This could be due to a high reactivity of cysteine residues which might have precluded high confidence peptide spectral matches during the analysis of LC-MS/MS data. Furthermore, mimicry between keratins and streptococcal M protein has been suggested to drive autoimmune responses in psoriasis in the past and T cell responses against keratin peptides in psoriasis patients have been demonstrated^28^. In our dataset, 41 keratin-derived peptides that are predicted to be HLA-C*06:02 binders were detected. Two of these peptides (SYLDRVRS**<u>L</u>** from KRT18 and S**<u>YL</u>**EKVRAL from KRT23 or KRT13) have associated streptococcal mimics: SYLQRVRS**<u>P</u>** from beta galactosidase and S**<u>LE</u>**EKVRAL from peptidoglycan-binding protein respectively, both are predicted weak binders of HLA-C*06:02. Another highly intriguing study has linked the SRGPVHHLL peptide with KIR-responses specific to HLA-C*06:02 and demonstrated strong engagement of this peptide-HLA-C*06:02 complex with KIR2DS1- and KIR2DL1-binding^29^. However, **<u>S</u>**R**<u>G</u>**PVHHL**<u>L</u>** is a synthetic peptide that could not be matched to a naturally occurring peptide. Interestingly, we found a similar peptide, **<u>L</u>**R**<u>P</u>**PVHHL, from the SUMO-interacting motif-containing protein 1 that has been detected in 1106 keratinocytes treated with IFN-γ that is also a strong binder of HLA-C*06:02. It would be of great interest to investigate whether any of these peptides are presented in patient lesional psoriatic skin and whether patients mount any CD8^+^ T cell, NK cell or NKT cell responses against these peptides.

To date, no *ex vivo* skin-derived ligands of HLA-C*06:02 have been reported. By generating the HLA-C06:02^Tg^ mouse, we were able to subject this new mouse model to the IMQ-induced model of psoriasis and confirm physiologic HLA-C*06:02 expression in lymphoid organs and skin. Skin inflammation scores in transgenic animals were comparable to the wildtype group. It needs to be noted that HLA-Cw6^Tg^ mice carry a fully human chimeric HLA-β2M monochain including the human α3 region which regulates physiologic expression levels while many other HLA transgenic mice use the murine α3 region in the construct to allow for HLA-restricted T cell responses. Thus, murine T cells in our model cannot recognise HLA-C*06:02-restricted antigens in the absence of human CD8, most likely the leading cause of similar PASI scores between wildtype and transgenic mice. Within the list of peptides detected exclusively in psoriasiform skin, structural source proteins like keratins, collagen, plakoglobin and vimentin are readily represented. Anti-vimentin responses have been implicated across several autoimmune diseases, most prominently in rheumatoid arthritis where a high proportion of patients have antibodies against citrullinated vimentin peptides^30^. Efforts to validate whether anti-vimentin antibodies play a role in psoriatic arthritis have yielded mixed results^31, 32^. However, considering that psoriasis is a T cell driven disease, HLA-C*06:02 restricted anti-vimentin CD8^+^ T cell responses might not necessarily translate to a humoral response. Indeed, the 8-mer RLDLERKV vimentin peptide, was unique to psoriasiform skin and exhibits all the hallmarks of an N-terminally truncated HLA-C*06:02 binder. Additionally, five further vimentin-derived peptides were found uniquely in psoriasiform skin, among them the strong binders ARLDLERKV and KRTLLIKTV. Among keratin peptides, AGFGSRSL from keratin 6A/B is of particular interest due to its predicted binding affinity to HLA-C*06:02 and a phosphorylated serine on position 5. Phosphorylation plays a central role in cell communication under inflammatory conditions, has been implicated in autoimmune disorders and we have previously shown that phosphopeptide MHC class I epitopes are stable thus enabling the generation of CD8^+^ T cell responses^33-35^. Intriguingly, keratin 6A is a hyperproliferation marker that is specifically expressed in patient lesional skin in contrast to healthy skin and previous reports have demonstrated that bacterial inflammation induces keratin 6A phosphorylation which in turn elevates cytoplasmic levels of the depolymerised filament^36, 37^. This hydrolysis leads to the formation of keratin-derived antimicrobial peptides termed KAMPs which have dual anti-bacterial and anti-inflammatory roles^38^.

Overall, the generated data of the skin-relevant HLA-C*06:02 immunopeptidome expands the known number of HLA-C*06:02 ligands by over six-fold. These experimentally validated immunopeptides can be used in future studies to explore pathogenic autoantigens in psoriasis. Further work into patient T cell responses and T cell cognate epitopes in psoriasis will be needed in order to establish broadly recognised antigen triggers of psoriasis. Such insights will allow studies into antigen-specific tolerance induction and develop novel therapeutic approaches with the prospect of inducing curative, long-lasting remission in psoriasis patients.

## Materials and Methods

### Cell line generation and culture

Full-length HLA-C*06:02 and HLA-B*57:01 were cloned into the pcDNA3.1 vector and stably transfected into A431 (A-431) cells by antibiotic selection. A431 keratinocytes and VMM1 melanocytes (CRL-3225) were maintained in 4.5g/L glucose DMEM supplemented with 10 % FCS, 2 mM glutamine, 1% (v/v) non-essential amino acids, 5 mM HEPES, 50 µM β-mercaptoethanol. 1106 keratinocytes (CCD 1106 KERTr or CRL-2309) were maintained in keratinocyte serum free medium (Gibco, 17005042) with addition of 0.05 mg/ml bovine pituitary extract and 35 ng/ml of EGF (Gibco, PHG0311). All cells received 100 IU/ml penicillin & 100 μg/ml streptomycin and were cultured at 37 °C/5 % CO2. In some conditions, cells were treated with human recombinant IFN-γ 1b (130-096-484, Miltenyi) at 2 ng/ml for 48h.

### Flow cytometry

Cell lines were stained with 5 μg DT9 or W6/32 (both from in-house preparation) followed by goat-α-mouse IgG – APC (17-4010-82, eBioscience). Mouse splenocytes were stained with Fc-block (553142 BD Pharmigen), live/dead aqua (423102, Biolegend), CD8-APC (17-0083-81, 17-0081-82, eBioscience), α-HLA-A/B/C-PE (311409, Biolegend, clone W6/32), α-HLA-A/B/C-PE-Cy7 (311430, Biolegend, clone W6/32), α-mouse-MHC I-PE (12-5998-82, eBioscience), α-human-B2M-FITC, α-mouse-b2m-BV605 (745120, BD Pharmigen), α-HLA-C-PE (clone DT9). For intracellular staining, cells were permeabilised with Saponin 0.3% before staining with DT9 or W6/32.

### Purification of peptide-HLA complexes

Cell pellets (0.5-0.8 × 10^9^ cells), supernatants (100-200 ml) and tissues were stored at -80°C until further use. Cell pellets and tissues were ground in a Retsch Mixer Mill MM 400 under cryogenic conditions; sHLA-containing supernatants were used directly. Following our standardised protocol^9^, cells and tissues were lysed in 0.5% IGEPAL (Sigma-Aldrich, #18896), 50 mM Tris, pH 8, 150 mM NaCl (Merck-Millipore, #106404) and protease inhibitors (Complete Protease Inhibitor Cocktail Tablet, 1 tablet per 50 ml solution; Roche Molecular Biochemicals, #11697498001) for 1 h at 4 °C with slow end-over-end mixing. Peptide-HLA complexes were immunoaffinity captured from clarified cell lysates (100,000 × g, 10 min) by passing through protein A sepharose resin which was either affinity-bound (mouse tissues) or additionally cross-linked to antibodies. The HLA-C specific DT9 antibody was used for cell lysates and supernatants, while mouse tissues were passed through pan-HLA class I antibody W6/32 bound resin. Lysates and supernatants were co-incubated with immunoaffinity beads for at least 1 hour. Bound and washed peptide-HLA complexes were eluted with 10% acetic acid. For tissues, the eluted mixture of peptides was purified by Amicon® 5kDa Ultra-Centrifugal filter unit (Merck Millipore) and concentrated by OMIX C18 Pipette Tips (Agilent, A57003100) prior to mass spectrometric analysis. For cells and supernatants, the peptide-HLA mixtures were fractionated off-line using a 4.6-mm × 100-mm monolithic reversed-phase C18 high-performance liquid chromatography (HPLC) column (Chromolith SpeedROD; Merck Millipore) and an ÄKTAmicro HPLC system (GE Healthcare). The mobile phase consisted of Buffer A (0.1% trifluoroacetic acid; Thermo Fisher Scientific) and buffer B (80% acetonitrile, 0.1% trifluoroacetic acid; Thermo Fisher Scientific). Peptide-HLA mixtures were loaded onto the column at a flow rate of 1 mL/min with separation based on a gradient of 2−40% Buffer B for 4 min, 40−45% for 4 min and a final rapid 2-min increase to 100%. Fractions (1 ml) were collected, pooled, vacuum-concentrated and diluted in 0.1% formic acid. Retention alignment peptide standards (iRT peptides^39^) were added to samples prior to mass spectrometric analysis.

### Mass spectrometry

Samples were analysed using a Q-Exactive plus mass spectrometer coupled online with a RSLC nano HPLC Ultimate 3000 (both Thermo Fisher Scientific). Samples were loaded on a 100 μm, 2 cm nanoviper pepmap100 trap column in 2% ACN, 0.1% formic acid at 15 μl/min. Peptides were eluted and separated at 300 μl/min on RSLC nanocolumn 75 μm x 50 cm, pepmap100 C18, 3 μm 100 Å pore size (Thermo Fisher Scientific), ACN increased from 2% to 10% over 1 min followed by a linear ACN gradient from 10% to 26% in 0.1% formic acid for 60 min followed by a linear increase to 34% ACN in 0.1% formic acid over 5 min and up to 80% ACN in 0.1% formic acid over 5 min, followed by decrease of ACN back to 2% and re-equilibration. The eluent was nebulised and ionised using the Thermo nano electrospray source with a distal coated fused silica emitter (New Objective, Woburn, MA, USA) with a capillary voltage of 1.7-2.2 kV. The instrument was operated in the data dependent mode automatically switching between full scan MS and MS/MS acquisition. Survey full scan MS spectra (m/z 375–1,800) were acquired in the Orbitrap with 70,000 resolution (m/z 200) after accumulation of ions to a 3 × 10^6^ target value with maximum injection time of 120 ms. Dynamic exclusion was set to 15 sec. The 12 most intense multiply charged ions (z ≥ 2) were sequentially fragmented by higher-energy collisional dissociation with a fixed injection time of 60 ms 17,500 resolution, AGC target of 1 × 10^4^ counts, 2 Da isolation width, underfill ratio 1% and 15 sec dynamic exclusion. Typical mass spectrometric conditions were as follows: spray voltage, 1.7 kV; no sheath and auxiliary gas flow; heated capillary temperature, 275 °C; normalised HCD collision energy 27%.

### Analysis of immunopeptidomics data

LC-MS/MS data from cell lines was searched against the human proteome using PEAKS Online 10 with a human database extracted from Uniprot 03/2022 and peptide identities subjected to strict bioinformatic criteria including the use of a decoy database to apply a false discovery rate (FDR) cut-off of 5%. The following search parameters were used: no cysteine alkylation, no enzyme digestion (considers all peptide bond cleavages), instrument-specific settings: Orbitrap, Orbi-Orbi (parent and fragment ion tolerance of 10 ppm and 0.02 Da respectively), HCD fragmentation, 8 to 25-mers, variable modifications set to: oxidation of methionine, N-terminal acetylation and deamidation of Asn/Gln. Additionally, Peaks PTM search was performed after a Peaks DB search with all default in-built modifications with same mass tolerance settings as Peaks DB. For the soluble HLA-C*06:02 immunopeptidome of A431_C6 cells, all peptides also detected in A431_B57 cells were excluded prior to any analysis. Gibbs clustering was performed with published available tools^40^.

Mouse immunopeptidomics data was searched with the same parameters but against the Uniprot C57BL/6 proteome appended with the human HLA-C*06:02 protein. All 8-12-mers were retained, peptides found in blank runs or wildtype C57BL/6 mice were excluded.

Cross-reactivity was assessed using the agrep UNIX command (https://github.com/PurcellLab/agrep_for_crossreactivity) to search for 1 or 2 mismatched amino acids between detected mammalian peptides and the Group A Streptococcal M6 proteome (UniProt 1167 download 2019_08).

### Generation of HLA-C*06:02 transgenic mice

HLA-C*06:02 is composed of three exons spanning 2.9 kb from ATG to the stop codon encoding for a 366 amino acid protein. β2m human gene is composed of three exons and extends over 7.4-kb (4.8-kb from ATG to stop). The tandem HLA-C*06:02 construct, composed of human B2M – linker - human HLA-C*06:02 mature cDNA - hGH polyA, was knocked in at the mouse b2m exon 2. A 45 amino acid linker developed by Lemonnier was used: (GGA GGT GGA GGT TCT) x 3. Genotyping was performed by Transnetyx.

### Imiquimod-induced model of psoriasis

Mice were kept under specific pathogen-free conditions and treated daily with a topical application of 62.5 mg of 5% imiquimod cream (Aldara™) on the left shaved and depilated flank and left ear while Vaseline was applied to the contralateral flank. Erythema, scaling and thickening were scored independently on a score of 0 to 5, from none to severe. The cumulative score of the three measured parameters provided the overall Psoriasis Area and Severity Index (PASI). All experiments were approved by the Monash Animal Research Committee MARP-2, AE17375.

### Histology

Fresh skin was frozen in OCT compound, cryosections were fixed with cold Acetone for 8 min and stained with DAPI to visualise nuclei.

### GOrilla pathway analysis

The combined immunopeptidome from 1106, A431_C6 and VMM1 cell lines was compared against published background data from B cell lines 721.221 and C1R and peptides detected in both datasets were removed^10, 16^. The resulting *in vitro* Epidermal Dataset was compared against 602 genes listed as skin-elevated in the Human Proteome Atlas, 118 of these genes were found in our data and were queried in GOrilla against the entire Epidermal Dataset^41^.

## Acknowledgments

We wish to thank the entire Monash Animal Research Laboratory team, FlowCore facility, Histology Platform and the Monash Proteomics & Metabolomics Facility.

